# Absolute Membrane Potential Recording with ASAP-Type Genetically Encoded Voltage Indicators Using Fluorescence Lifetime Imaging

**DOI:** 10.1101/2025.08.08.669310

**Authors:** Anagha Gopalakrishnan Nair, Marko Rodewald, Hyeonsoo Bae, Philipp Rühl, Jürgen Popp, Michael Schmitt, Tobias Meyer-Zedler, Stefan H. Heinemann

## Abstract

The electrical membrane voltage (*V*_m_) characterizes the functional state of biological cells, thus requiring precise, non-invasive *V*_m_-sensing techniques. While voltage-dependent fluorescence intensity changes from genetically encoded voltage indicators (GEVIs) indicate *V*_m_ changes, variability in sensor expression confound determination of absolute *V*_m_. Fluorescence lifetime imaging microscopy (FLIM) promises a solution to this problem, as fluorescence lifetime is expected to be unaffected by sensor expression and excitation intensity. By examining ASAP1, ASAP3, JEDI-1P, rEstus, and rEstus-NI (G138N:T141I) with one-photon excited FLIM measurements, we demonstrate that all sensors display a voltage-dependent lifetime. With the highest lifetime change in the *V*_m_ range of -100 to 50 mV of about 730 ps, ASAP3 and rEstus-NI are preferred for FLIM recordings. At a physiologically relevant *V*_m_ of -30 mV, the voltage sensitivity of rEstus-NI (6.6 ps/mV) is 3.6 and 1.4 times greater than that of ASAP1 and rEstus, respectively. As a proof of concept, we successfully used rEstus-NI to estimate absolute resting *V*_m_ in HEK293T, A375 melanoma, and MCF7 breast cancer cells and quantified spontaneous *V*_m_ fluctuations in A375 cells.

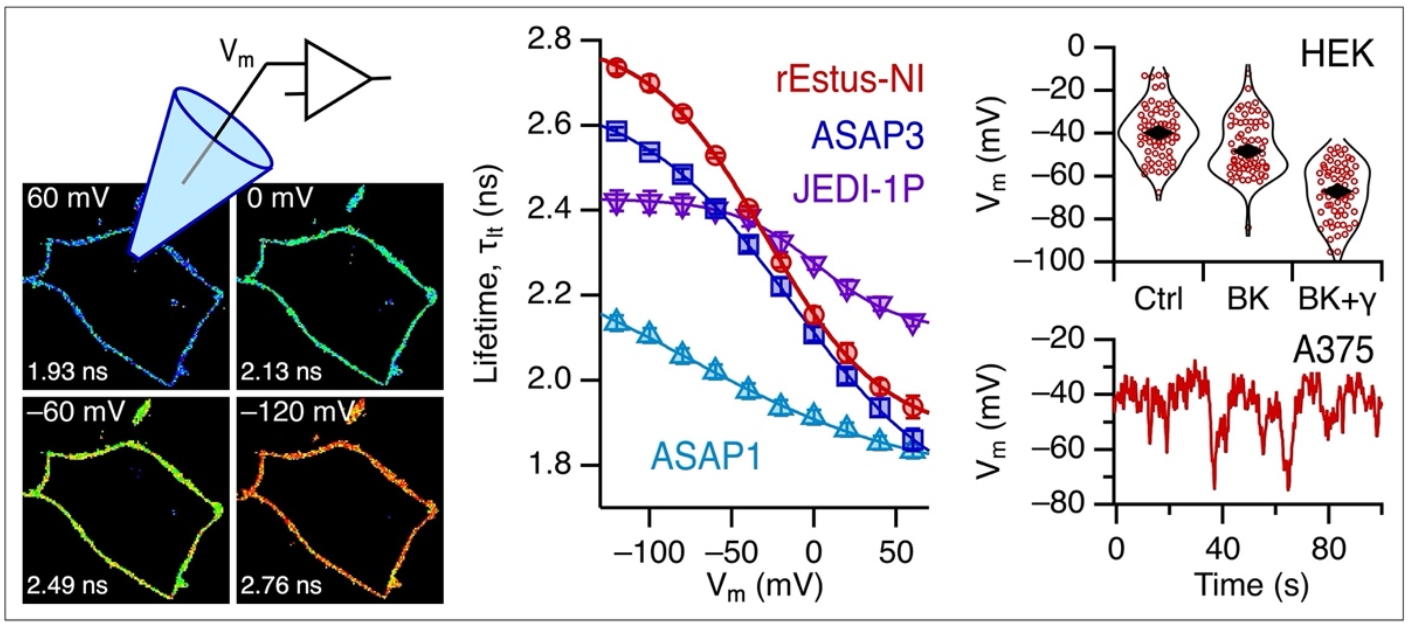

## INTRODUCTION

The electrical membrane potential (*V*_m_) is a key physiological property of biological cells. During action potentials in neurons and muscle fibers, *V*_m_ undergoes about 100 mV changes within milliseconds. *V*_m_ changes of much smaller amplitude, occurring over seconds to hours, accompany essential processes such as cell-cycle progression ^1-3^, cancer cell growth and migration ^4, 5^, tissue formation, and wound healing ^6, 7^. The causal relationship between *V*_m_ changes and the aforementioned physiological and pathophysiological functions remains a matter of intensive research. Non-invasive sensors to accurately measure *V*_m_ are therefore vital for gaining a deeper understanding of the underlying biological and physiological mechanisms.

Electrophysiological techniques are regarded as the gold standard for measuring *V*_m_ but suffer from low throughput and high invasiveness. Optical methods employing voltage-sensitive dyes or genetically encoded voltage indicators (GEVIs) provide an alternative, offering spatial resolution, reduced invasiveness, ease of use, and high throughput ^8^. While voltage-sensitive dyes encounter challenges related to complex loading procedures, internalization, and interference with membrane proteins ^9, 10^, GEVIs address some of these drawbacks and serve as powerful tools for *V*_m_ measurement^11,12^.

Fluorescence intensity (*F*) and fluorescence lifetime (τ_lt_) are the two most commonly used voltage-dependent factors for bioimaging of voltage-dependent chromophores. While intensity-based *V*_m_ measurements can detect fast action potentials and subthreshold events, they cannot provide absolute *V*_m_ values ^13, 14^. Single-color fluorescent intensity recording fails to measure absolute *V*_m_ due to signal variability caused by sensor expression, cell morphology, photobleaching, and variations in light intensity ^15^. Even with individual calibration for each instrument, absolute *V*_m_ measurements are problematic ^15, 16^. While ratio-based imaging strategies can correct for concentration-dependent fluorescence changes ^17, 18^, wavelength-dependent photobleaching can introduce errors ^18^. Analyzing the ratio of photon emission from two fluorophores within the same sensor can eliminate motion-induced optical signal changes ^19, 20^. The fusion of a second voltage-independent protein allows for absolute *V*_m_ measurements, but incorporating a second protein can affect membrane trafficking, reduce sensor expression levels, and may be impacted by unequal pH sensitivity of the chromophores^12^.

Fluorescence lifetime-based measurements of *V*_m_ promise improved quantification for optical assessments of absolute *V*_m_. Fluorescence lifetime imaging microscopy (FLIM) measures the duration a molecule remains in an excited state before returning to a lower energy state by emission of a photon, a parameter that strongly depends on the chromophore’s molecular environment. Unlike fluorescence intensity, in a first approximation fluorescence lifetime is independent of the fluorophore concentration, bleaching, excitation intensity, and the spectral properties of the detection system ^21, 22^. In principle, this makes FLIM advantageous compared to intensity-based measurements, and attempts to use FLIM with GEVIs, voltage-sensitive dyes, and genetically encoded Ca^2+^ indicators to report absolute *V*_m_ or the intracellular Ca^2+^ concentration have shown some promise ^16, 23-26^.

In GEVIs employing Förster resonance energy transfer (FRET), changes in τ_lt_ arise from variations in FRET efficiency^23^. However, GEVIs exhibiting the highest voltage sensitivities of *F* are based on voltage-sensing domains (VSDs), with the ASAP family of sensors utilizing a VSD fused to a circularly permuted GFP (cpGFP) (Fig. S2a). As these VSD-based sensors do not rely on quenching of a FRET donor, the relationship between *F* and τ_lt_ is non-trivial. Changes in *F* can result from alterations in the absorption coefficient or the fluorescence quantum yield, while only the latter directly affects the fluorescence lifetime^27^. A previous study has demonstrated voltage-dependent changes in τ_lt_ of ASAP1 ^28^ under two-photon excitation^23^.

Here we examined the voltage-dependent fluorescence lifetime of the state-of-the-art ASAP1-derived sensors ASAP3 ^29^, rEstus^12^, and JEDI-1P ^30^. We found that ASAP3 and rEstus exhibit substantially higher voltage-dependent lifetime changes compared to ASAP1 and JEDI-1P. For all ASAP derivatives, the voltage of maximal sensitivity for lifetime changes (*V*_hτ_) was shifted to more positive values compared to the voltage of maximal sensitivity for intensity changes (*V*_hF_). We thus selected a rEstus variant (rEstus-NI) for which the voltage of highest sensitivity of τ_lt_ changes is aligned with physiologically relevant resting membrane potentials. rEstus-NI enabled absolute *V*_m_ measurements of steady-state *V*_rest_ in cancer cell lines, as well as the detection of spontaneous, dynamic *V*_m_ changes in non-excitable cells.

## EXPERIMENTAL SECTION

### Generation of DNA plasmids for expression in mammalian cell lines

Expression plasmids of the following genetically encoded voltage indicators were constructed using standard molecular biology methods: ASAP1 ^28^, ASAP3 ^29^, rEstus ^12^, and JEDI-1P ^30^. rEstus was derived from ASAP3, harboring the mutations N138G, Y141T, and Q396R in the voltage-sensing domain and S318A in the cpGFP part of the sensor (rEstus = ASAP3-N138G:Y141T:S318A:Q396R). During the optimization for FLIM applications, the following variant was selected from a previous mutagenesis screen^12^: rEstus-G138N:T141I (rEstus-NI), which is equivalent to ASAP3-Y141I:S318A:Q396R. All mutation positions are given relative to the N terminus of ASAP3. To determine the relative molecular brightness, all GEVI constructs were also produced as fusion constructs with the red fluorescent protein mKate2 being placed at the N terminus (thus intracellular) of the GEVI, as previously described ^11^.

To study the impact of K^+^ channel expression on *V*_m_, we used the human large-conductance Ca^2+^- and depolarization-activated K^+^ channel α subunit (*KCNMA1*, BKα, NP_002238). LRRC26 (NP_001013675), a leucine-rich repeat-containing protein that functions as an auxiliary γ1 subunit of BK_Ca_ channels (BKγ1), was fused to the C-terminus of BKα with a short linker (encoding the sequence “KLT”) using a *Hind*III site. The BKα and BKγ1 proteins form a complex after cleavage by endogenous peptidases at an endogenous signal peptide within LRRC26, as described previously ^31^.

All variants were inserted into a pcDNA3.1 vector with a CMV promoter. Cloning primers were synthesized using Sigma-Aldrich’s DNA oligos service. All constructs were verified by DNA sequencing.

### Cell culture and transfection

Human embryonic kidney 293T (HEK293T, CAMR; Porton Down, Salisbury, UK) cells and MCF-7 cells (ECACC; Porton Down, Salisbury, UK) were cultured in a medium consisting of a 1:1 mixture of Dulbecco’s Modified Eagle’s Medium and Nutrient Mixture F-12 (DMEM-F12, Thermo Fisher Scientific, Waltham, MA, USA), supplemented with 10% fetal bovine serum (FBS). The cells were maintained in a humidified incubator at 37°C with 5% CO_2_. A375 cells (ATCC, Manassas, VA, USA) were cultured in DMEM (Sigma Aldrich) supplemented with 10% FBS, at 37°C with 10% CO_2_.

For combined electrophysiology and FLIM experiments, cells were plated on 35-mm glass-bottom dishes (Ibidi, Martinsried, Germany) at a density of 10,000 cells per dish. HEK293T cells were transfected with 1 µg of plasmid DNA per dish using the ROTI®Fect transfection kit (Carl Roth, Karlsruhe, Germany) the following day. MCF-7 and A375 cells were transfected with 1 µg of DNA using the SF Cell Line 4D-Nucleofector kit (Lonza, Basel, Switzerland) according to the manufacturer’s instructions by electroporation (4D-Nucleofector, Lonza).

For FLIM imaging experiments, 0.5 µg of *KCNMA1* or *KCNMA1*-*LRRC26* plasmid was co-transfected with 0.5 µg of rEstus-NI DNA per dish. To minimize cell-cell contacts, cells were trypsinized and reseeded one day after transfection. The actual recordings were obtained two days post-transfection.

In indicated cases, cells were fixed with 1 mL of 4% paraformaldehyde (PFA) for 5 min. After removing the PFA, the cells were washed with 1 mL PBS and subsequently kept in 2 mL PBS for recording.

### Electrophysiological control of the membrane voltage

The fluorescent voltage sensors were calibrated using simultaneous fluorescence and fluorescence lifetime measurements and electrophysiological patch-clamp recordings. Whole-cell patch-clamp measurements were conducted with an EPC10 amplifier and PatchMaster acquisition software (HEKA Elektronik, Lambrecht, Germany). Pipettes were fabricated from borosilicate glass with filament; tips were coated with dental wax and fire-polished to yield resistances between 1.0 and 2.0 MΩ. The series resistance was corrected electronically by up to 75%. The holding membrane voltage was -60 mV, and membrane voltages ranging from -120 mV to 60 mV in steps of 20 mV were examined, each step lasting 4 s, during which 42 FLIM frames were acquired (Fig. 1a, *top*). The first segment at -60 mV was longer than the subsequent segments in order to saturate the loss of fluorescence intensity caused by photoswitching.

**Figure 1.**
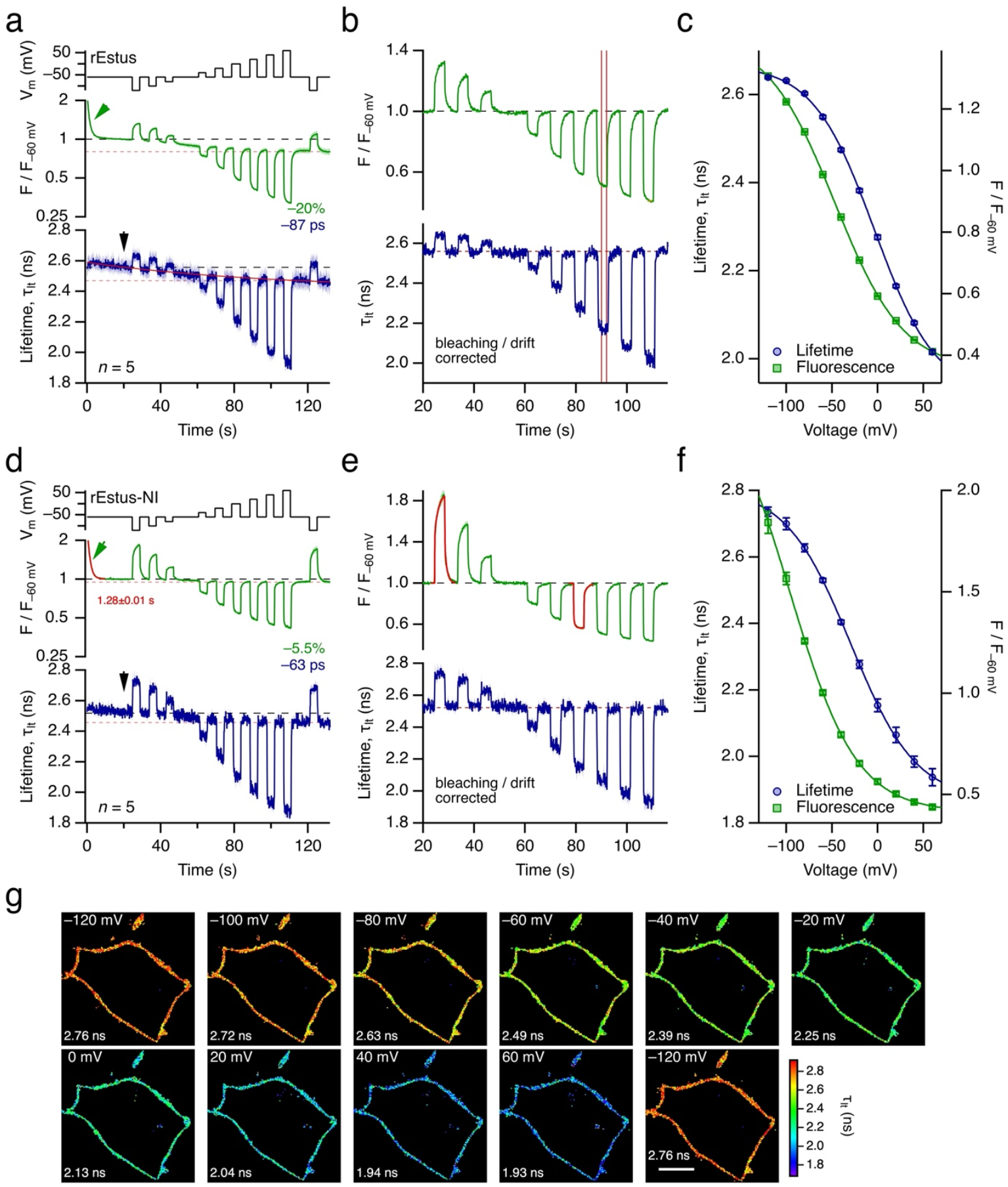
Fluorescence lifetime recordings of rEstus (a-c) and rEstus-NI (d-g) in HEK293T cells under patch-clamp control. (**a**) Voltage profile (*top*), fluorescence normalized to the fluorescence at -60 mV after 20 s of recording time (*F*/*F*_-60 mV_, *middle*) on a logarithmic scale, and fluorescence lifetime (_lt_, *bottom*). *F* and _lt_ are mean values of 5 independent cells under voltage-clamp control; SEM indicated by shading. The green arrow indicates the loss of fluorescence at the beginning of the recording, presumably as a result of photoswitching. Estimates of time-dependent drifts, between 20 s and 130 s, in fluorescence (bleaching) and fluorescence lifetime (red dashed lines) are also indicated. The continuous red line in the _lt_ plot indicates the exponential approximation of the time-dependent drift in fluorescence lifetime. The arrow marks the reference time for drift correction. (**b**) As in (a) but after exponential correction of the time-dependent drifts; *F*/*F*_-60 mV_ is shown on a linear scale. Vertical lines indicate the integration range of one voltage pulse. (**c**) Mean fluorescence lifetime (blue) and normalized fluorescence intensity (green) of the data shown in (b) as a function of voltage. The superimposed curves are data fits according to Eq. 1, yielding: *F*_max_ = 1.444, *F* = 1.092, *V*_hF_ = -47.1 mV, *k*_hF_ = 37.8 mV; _max_ = 2.668 ns, = 0.750 ns, *V*_h_ = -3.5 mV, *k*_h_ = 33.7 mV. (**d**) As in (a) for rEstus-NI. (**e**) As in (b) for rEstus-NI. To estimate the kinetics of the change in fluorescence following step-wise voltage changes, resulting fit curves are superimposed for the recordings at -120 and 0 mV (for full analysis, see Fig. S3). (**f**) As in (c) for rEstus-NI. The superimposed curves are data fits according to Eq. 1, yielding: *F*_max_ = 2.60, *F* = 2.19, *V*_hF_ = -96.6 mV, *k*_hF_ = 36.6 mV; _max_ = 2.81 ns, = 0.94 ns, *V*_h_ = -29.4 mV, *k*_h_ = 35.8 mV. (**g**) FLIM images of a single rEstus-NI-expressing HEK293T cell under whole-cell patch-clamp control at the indicated voltages. Scale bar: 10 µm. The mean lifetimes are indicated.

The bath solution was (in mM) 146 NaCl, 4 KCl, 2 CaCl_2_, 2 MgCl_2_, and 10 HEPES; pH 7.4 (NaOH). The pipette solution contained (in mM) 130 KCl, 2.5 MgCl_2_, 10 EGTA, and 10 HEPES; pH 7.4 (KOH). For the indicated experiments, the concentration of KCl in the extracellular recording buffer was reduced to 2 mM. All recordings were obtained at room temperature (20-23°C).

For FLIM measurements of cells without voltage-clamp control, cell culture dishes were prepared in the same manner as for patch-clamp experiments. A375 and MCF-7 cells were transiently transfected with DNA coding for rEstus-NI. Before the experiment, the transfected cells were washed with 1 mL of external solution containing 5 mM glucose, and then kept in 2 mL of the same solution. For experiments involving gramicidin to short-circuit the membrane voltage, 1 µL of a 2 mM stock solution of gramicidin D (G5002, Sigma Aldrich) in ethanol was added to 2 mL of external solution with 5 mM glucose. Imaging was performed 10-20 min after the medium was exchanged.

### Fluorescence lifetime imaging

Fluorescence lifetime imaging experiments were conducted using an inverted confocal laser-scanning microscope equipped with a white-light laser (DMI8 inverted microscope and SP8 Falcon confocal laser scanner, Leica, Germany). Data were acquired with Leica Application Suite X version 3.5.7 software. To interface confocal FLIM imaging with patch-clamp electrophysiology, the live data mode software module and trigger box were employed. The white-light laser (NKT Photonics, Denmark) was operated at 85% power; 20% of the laser power at 480 nm was used for excitation, with a repetition rate of 40 MHz. Full laser power at 480 nm corresponds to 20 µW of average laser power in the focal plane of the 63x oil immersion microscope objective (HC Plan Apo CS2, Leica). Fluorescence lifetime images were measured with a confocal hybrid detector (HyD SMD, Leica) by time-correlated single-photon counting in the spectral range of 490 to 600 nm in xyt scanning mode using the following scanning parameters: zoom 5, 36.9 µm field of view, 128 × 128 pixels, scan speed of 1400 lines/s, and an 88.12 µm pinhole size corresponding to 1 airy unit at 535 nm. Lifetime data acquisition was performed with a 100 ps sampling interval and a 25 ns time window, thus collecting 250 temporal data points.

For calibration measurement of voltage-clamped cells, 1,350 images were recorded, resulting in a total acquisition time of about 2 min. The frame rate of 10.29 frames per second allowed for the acquisition of dynamic changes on a time scale of 100 ms.

Data analysis was performed using the LAS X FLIM/FCS software (Version 3.5.6, Leica). A region of interest was manually drawn to select the cell membrane; it was verified that the cells did not move out of the ROI during the recording. The individual images were analyzed using the all-photon filter, and the lifetime was fitted using the n-exponential deconvolution function with two exponential components and five fitting parameters (two lifetimes, two amplitudes, and an offset). The time interval for analysis was adjusted to exclude reflection peaks. Intensity-weighted mean lifetimes (τ_lt_) were used as the resulting parameter to limit the errors introduced by the fast lifetime component, which was close to the resolution limit of the photon counting system.

For FLIM imaging experiments without patch-clamp control, fluorescence lifetime was measured using the same excitation and emission settings as described earlier. The scanning parameters included a zoom factor of 2, a field of view measuring 92.26 µm, a resolution of 128 × 128 pixels, a scan speed of 700 lines per second, and a pinhole size corresponding to 1 airy unit at 535 nm. Lifetime data acquisition was carried out with about 10 frames/s. For each field of view, 400 images were recorded. To mitigate the effects of photoswitching, the median lifetime values were calculated using only the second half of the total images obtained from cells under various conditions. The corresponding voltage for each median lifetime was subsequently determined using the τ_lt_-V calibration curve for each cell line. For image processing, see Suppl. Material.

### Analysis of voltage dependence

In GEVI calibration experiments, mean fluorescence (*F*) and intensity-weighted lifetimes (τ_lt_) for the respective ROI were measured as a function of time using the voltage-clamp protocol shown in Fig. 1a. Fluorescence was normalized to the *F* values obtained in the first -60 mV segment. Drifts in *F* (mostly due to bleaching) and τ_lt_ over time were corrected by fitting single-exponential functions to the time courses at -60 mV and by subtracting the fit function relative to the end of the first segment at -60 mV.

For the compilation of fluorescence–voltage (*F*-*V*) and fluorescence lifetime–voltage (τ_lt_-*V*) relationships under voltage-clamp control, drift-corrected *F* and intensity-weighted τ_lt_ values during the last 20 frames (corresponding to about 2 s) at the respective voltage were averaged and plotted as a function of voltage (*V*_m_). The voltage dependence was described with a Boltzmann distribution, characterized by a value at saturating voltage (*F*_∞_ or τ_∞_), the maximal voltage-dependent change (Δ*F*,Δτ), the voltage of half-maximal change (*V*_hF_, *V*_hτ_), and the steepness factor (*k*_hF_, *k*_hτ_). For the example of the voltage dependence of fluorescent lifetime:

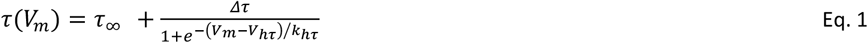

Final data analysis and figure generation were performed with Igor Pro 9 software (WaveMetrics, Lake Oswego, OR, USA).

### Statistical analysis

Data are presented as means ± SEM, unless specified otherwise. The number of individual measurements is denoted as *n*. Statistical tests are given in the text.

## RESULTS AND DISCUSSION

### Fluorescence lifetime imaging of *V*_m_ using ASAP1-derived GEVIs

In 2015, Brinks *et al*. showed that τ_lt_ of ASAP1 depends on voltage ^23^. Since then, several ASAP1-derived GEVIs with improved properties, such as increased sensitivity, speed, and brightness, have been developed. Given the brightness and high voltage sensitivity of *F* at *V*_rest_ of rEstus ^12^, we examined the usefulness of this sensor and derivatives for τ_lt_ recordings for absolute *V*_m_ determination.

After expression of the GEVIs in HEK293T cells, *F* and τ_lt_ were measured using a confocal laser-scanning microscope equipped with a white-light laser for excitation. *V*_m_ of the cells was controlled with whole-cell patch-clamp. For further analysis, we used the membrane-delimited *F* and the pixel-wise measured τ_lt_ (Fig. S1a). The latter was determined by fitting the arrival time-distribution of photon counts after an excitation pulse with deconvoluted double-exponential functions (Fig. S1b), then yielding the intensity-weighted average τ_lt_. Images were taken at a frame rate of about 1/100 ms while the voltage followed the protocol shown in Fig. 1a (*top*). The time course of mean *F* and τ_lt_ values for rEstus are shown in Fig. 1a. As indicated earlier ^12^, *F* is strongly voltage-dependent, with a change by about 70% when the voltage is altered from -120 to 60 mV. As is typical for cpGFP variants, there is an initial fluorescence loss, presumably due to photoswitching ^32^. Under the illumination paradigm used for FLIM applications, the time constant for photoswitching was about 1 s, thus becoming saturated before the calibration protocol began. During the course of this calibration experiment (132 s), *F* at -60 mV diminished by about 20%, presumably due to photobleaching. The mean τ_lt_ exhibited a strong dependence on voltage, ranging from 2.64 ns at -120 mV to 2.02 ns at 60 mV. Notably, τ_lt_ was not affected by the prominent photoswitching at the start of the recording. τ_lt_ values at -60 mV decreased by 87 ps during the course of the calibration experiment. The time-dependent changes in *F* and τ_lt_ were corrected for estimated exponential bleaching (*F*) and drift (τ_lt_), thereby considering the data values at 20 s following image acquisition (black arrow in Fig. 1a, *bottom*) as a reference (Fig. 1b).

*F* and τ_lt_ were averaged for the second half of the 4-s voltage pulses to account for the initial rising phase, which was particularly prominent in *F*(*t*). The voltage dependencies of the resulting mean values (Fig. 1c) were described using Boltzmann distributions (Eq. 1) to yield the maximal voltage-dependent signal change, as well as the half-maximal voltage (*V*_h_) and the slope factor (*k*_h_) characterizing the voltage dependence. The following parameters characterize rEstus as a FLIM-based sensor for *V*_m_: there is a change in τ_lt_ (Δτ_lt_) of 620 ps between -120 and 60 mV, and τ_lt_ is not affected by photoswitching. *V*_h_ values derived from *F* (*V*_hF_) and τ_lt_ (*V*_h τ_) differ by about 45 mV (−47.1 vs. -3.5 mV), while the voltage dependencies are about the same (*k*_hF_ 37.8 and *k*_h τ_ 33.7 mV, respectively).

The difference between the optimal voltage sensitivity of *F* and τ_lt_ places *V*_hτ_ near complete cell depolarization. For studying typical resting membrane voltages, a GEVI with a left-shifted *V*_hτ_ would be desirable. Therefore, we evaluated rEstus variants originating from site saturation mutagenesis of residues G138 and T141^12^, located in segment 3 of the voltage-sensing domain (Fig. S2a), and selected G138N:T141I (rEstus-NI or ASAP3:S318A:Y141I:Q396R) for further analysis in FLIM experiments. As shown in Fig. 1d and e, Δτ_lt_ of rEstus-NI was even larger than that for rEstus: 798 ps between -120 and 60 mV. Similar to rEstus, τ_lt_ is not affected by photoswitching. Within the calibration of 132 s, *F* at -60 mV diminished by about 5.5% and τ_lt_ values at -60 mV decreased by 63 ps. *V*_h_ values derived from *F* (*V*_hF_) and τ_lt_ (*V*_hτ_) differ by about 66 mV (−95.4 vs. -29.2 mV), while the voltage dependencies are about the same (*k*_hF_ 36.6 and *k*_hτ_ 35.9 mV, respectively) (Fig. 1f). Thus, the mid-point of the τ_lt_-V relationship of rEstus-NI is left-shifted by about 25 mV with regard to rEstus and therefore better suitable for physiological applications.

To compare the lifetime-dependent signal of rEstus and rEstus-NI with other state-of-the-art sensors, we also measured the voltage-dependent τ_lt_ of ASAP1 and its more recent derivatives, ASAP3, and JEDI-1P. We confirmed the voltage dependence of τ_lt_ for ASAP1, as previously shown by Brink and colleagues ^23^: Δτ_lt_ was 310 ps, varying from 2.14 ns to 1.83 ns (Fig. 2a-b). This change is less than half of the Δτ_lt_ of rEstus-NI. Estimates based on fitting a Boltzmann distribution to τ_lt_(*V*) (Eq. 1) lead to the conclusion that ASAP1 is not suitable for FLIM measurements due to the very negative half-maximal voltage *V*_h τ_ of -73.7 mV, thus leaving most of the voltage-dependent Δτ_lt_ outside the physiological *V*_m_ range.

**Figure 2.**
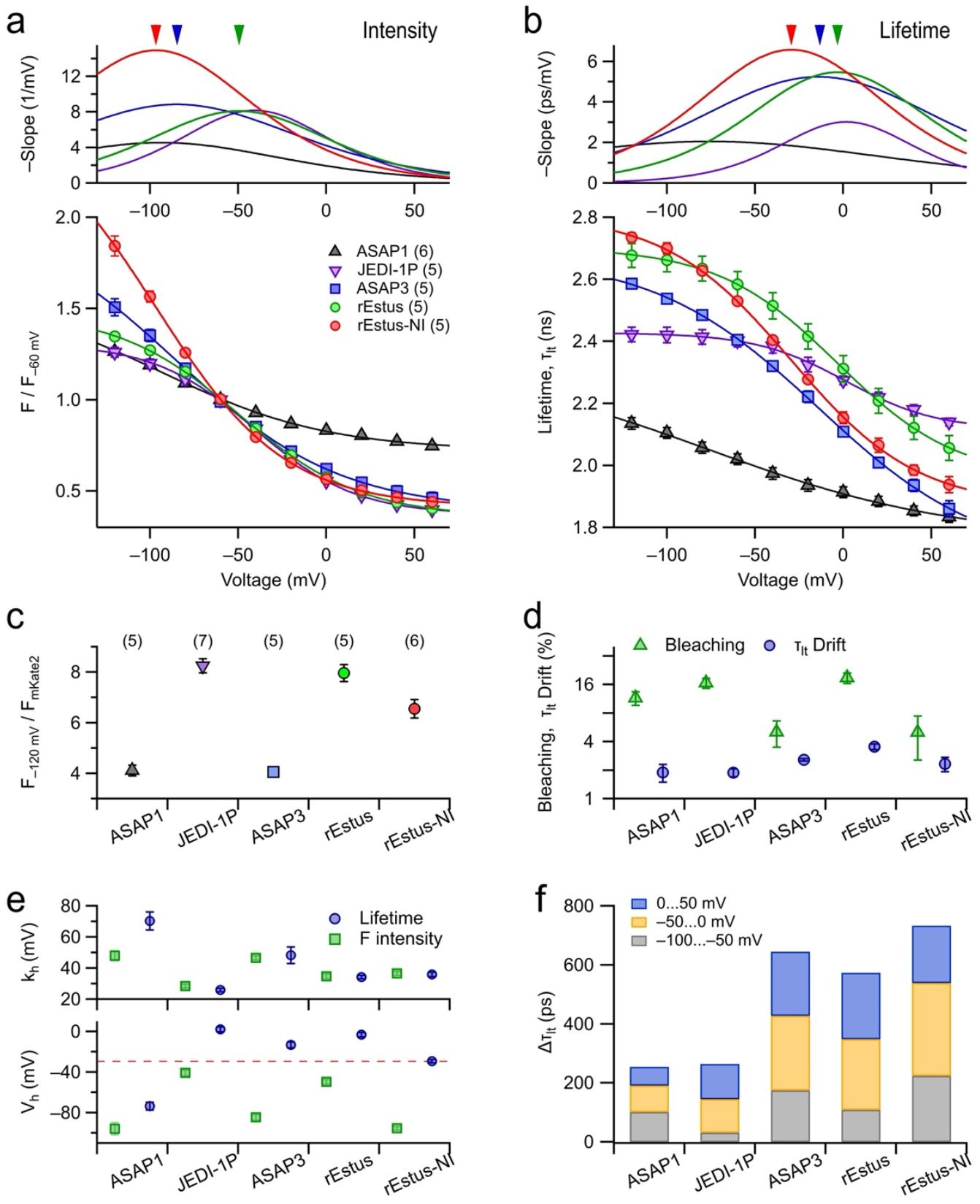
Comparison of GEVIs from the ASAP family. Experiments, as described for rEstus and rEstus-NI in Fig. 1, were conducted for the GEVIs ASAP1, JEDI-1P, and ASAP3. (**a**) *bottom*: Mean fluorescence intensity, normalized to -60 mV, as a function of voltage for the indicated GEVIs, with superimposed fits according to Eq. 1. Error bars indicate ± SEM, and the number of cells measured is given in parentheses. *top*: First derivative of the fit functions to the *F*-*V* data, indicating the voltage sensitivity. The arrowheads mark the voltage of maximal sensitivity for ASAP3 (blue), rEstus (green), and rEstus-NI (red). (**b**) Analysis and symbols used as in (a) for fluorescent lifetime data. (**c**) Mean molecular brightness (Suppl. Methods) at -120 mV, relative to the brightness of mKate2. For the full *F*-*V* relationships, see Fig. S2. (**d**) Estimated fluorescence bleaching (triangles) and relative drift in fluorescence lifetime (circles) during the 130-s period of repetitive image acquisition as needed for the calibration for the indicated GEVIs. (**e**) Parameters characterizing the voltage dependence of *F* (green) and _lt_ (blue) from the data fits shown in (a) and (b), respectively. (**f**) Cumulative change in fluorescence lifetime for the indicated voltage ranges, derived from the data fits in (b).

ASAP3 exhibited a total Δτ_lt_ of 726 ps with *V*_hτ_ of -13.2 mV – a mid-point voltage between that of rEstus and rEstus-NI. JEDI-1P performed less favorably in FLIM applications; its τ_lt_ ranged from 2.42 ns (−120 mV) to 2.14 ns (60 mV), resulting in a total Δτ_lt_ of only 280 ps, which is even smaller than that of ASAP1. *V*_hτ_ of JEDI-1P was 2.1 mV, indicating a substantial rightward shift compared to ASAP1. It also showed an approximate 43-mV shift relative to its own *V*_hF_. Of all tested sensors, rEstus-NI and ASAP3 performed best in FLIM recordings, with the largest lifetime change and a well-positioned *V*_hτ_; they were followed by rEstus, ASAP1, and JEDI-1P (Fig. 2f).

Since the sensor brightness is an important factor for the accuracy of τ_lt_ determination, we estimated the molecular brightness of the voltage-sensitive cpGFP moiety of the sensors by generating N-terminal fusion proteins with voltage-independent red-fluorescent mKate2, which compensates for variations in expression levels ^12^. While ASAP3 and rEstus-NI provide a large Δτ_lt_ within the physiological voltage range, rEstus and JEDI-1P are approximately twice as bright as ASAP3 and ASAP1 (Fig. 2c, Fig. S2); the brightness of rEstus-NI is intermediate.

The Δτ_lt_ of about 250 ps for ASAP1 ^28^ between -100 and 50 mV aligns well with results from earlier two-photon FLIM measurements ^23^. The sensitivity of ASAP1 at -50 mV of less than 2 ps/mV defines clear limits of this GEVI for FLIM-based *V*_m_ measurements. The same applies to JEDI-1P ^30^ (1 ps/mV). ASAP3 ^29^ and rEstus ^12^, both very efficient *F*-based *V*_m_ indicators, performed much better in FLIM with maximum sensitivities of about 5 ps/mV. rEstus-NI, with a peak sensitivity of about 6 ps at -50 mV and a Δτ_lt_ of 730 ps, ranges on top of the ASAP family.

In addition to the absolute voltage-dependent changes in *F* or τ_lt_, the voltage of the largest sensitivity (*V*_h_) and the steepness factor (*k*_h_) are important parameters for sensor optimization. For all GEVIs investigated, *V*_hτ_ values (blue squares) are right-shifted with respect to the *V*_hF_ values (green circles), while the voltage sensitivities (*k*_h_) of *F*(*V*) and τ_lt_(*V*) are approximately the same (Fig. 2e; ASAP1 appears to be an exception, but for this sensor *k*_h_ cannot be estimated well because of its strongly left-shifted *V*_h_.). This difference could partly be related to an unequal voltage dependence of the quantum yield and the absorption coefficient ^27, 33^. Based solely on *F*(*V*), it cannot be predicted if a cpGFP-based GEVI is suitable for FLIM-based *V*_m_ measurements.

Conformational changes in the VSD may alter the absorption coefficient and/or the lifetime of the excited state. We observed a slow component in the *F* traces, which was predominantly seen at negative voltages and was absent in the lifetime recordings. It has been shown that the VSD of voltage-sensing phosphatases resides in the down or down-minus state at very negative voltages, while at positive voltages, it makes a transition to the up or up-plus state ^34^. As demonstrated for rEstus-NI in Fig. 1, the fraction of *F* related to lifetime-dependent changes was larger at positive voltages and strongly diminished at negative voltages. This suggests that during a depolarizing voltage step the initial section of the VSD movement from the bright down state primarily affects the absorption coefficient – potentially through recovery from reversible cis-trans isomerization of the chromophore and/or a change in the degree of the protonation of the chromophore – while the later part of the movement predominantly influences the excited-state lifetime.

The initial photoswitching of rEstus-NI (Fig. 1d) was similar to that of rEstus (Fig. 1a); the slow loss of *F* during a calibration measurement, presumably resulting from photobleaching, was only 5.5%, and the drift in τ_lt_ at -60 mV amounted to -63 ps. While a certain drift in τ_lt_ was observed for all examined GEVIs, ASAP3 and rEstus-NI appear better than the others with respect to bleaching (Fig. 2d). In experiments in which τ_lt_ was only measured at the begin and the end of the 130-s episode without any light application between, the drift in τ_lt_ was diminished to -9.8 ± 2.2 ps (*n* = 9), indicating that the drift is primarily induced by photochemical modifications and not merely by the recording time. Perhaps there is a slight bias arising from a differential trend in the amplitudes of the fast and slow fluorescence lifetime components. Whatever the reason for the minor decline in τ_lt_, it does not present a serious problem if calibration and final measurements are conducted within a similar time frame and illumination intensity.

The kinetics of *F* and τ_lt_ following a sudden voltage step was markedly different. For voltage steps starting from -60 mV (termed “on”), there is a prominent slow phase in *F* with a time constant, τ_s,on_, of about 1.5 s with a relative fraction (*r*_s_) of 45% at -120 mV (Fig. 1e; for full analysis, see Fig. S3); however, the fluorescence lifetime does not show this component, rendering its time course limited by the time resolution of the FLIM measurements. The slow components in *F* during the repolarizing voltage steps to -60 mV (τ_s,off_) were also partially observed in the time course of τ_lt_ (Fig. 1e). Thus, only part of the voltage-dependent fluorescence emitted by rEstus-NI is directly related to *F* changes resulting from lifetime changes. The amplitude of this fraction exhibits an exponential voltage dependence, tending to about 18% at strongly depolarized membranes (Fig. S3). The voltage dependencies of the fast component and the maximum amplitude in *F*(*t*) after voltage changes (Fig. S3) suggest that the mid-voltage of τ_lt_(*V*) (*V*_hτ_ = -29.2 ± 1.4 mV) appears to be related to *V*_hF_ of the fast component (−56.0 ± 2.3 mV) (*V*_hF_ of the maximal fluorescence was -93.4 ± 2.9 mV). The slow component is not seen in τ_lt_ and, therefore, may result from equilibration of the fluorophore’s photoswitching.

In principle, ASAP-based GEVIs can be used ratiometrically by exciting at 400 and 480 nm ^12^. However, the GEVI’s fluorescence at 400 nm is not only small but also typically confounded by endogenous background fluorescence, such that recording the lifetime with excitation at 480 nm offers advantages. Moreover, τ_lt_ does not seem to be affected by photoswitching. Since *F* is diminished by about 30% after the start of illumination, pre-illumination of more than 1 s is needed for the fluorophore to reach an equilibrium in the photoswitched state before *F* data can be collected. This initial transient component is absent in the time course of τ_lt_. In addition, while step-changes in *V*_m_ result in a fast and slow *F* response, τ_lt_ is dominated by a fast component. Although the underlying mechanisms remain to be investigated, the primary reason appears to be a voltage dependence of the photoswitching equilibrium, which affects *F* but which does not impact τ_lt_. The absence of a slow component, as seen in the *F*(*t*) traces, is an advantage when recording τ_lt_(*t*) in FLIM experiments, thus providing a stable *V*_m_ signal about 200 ms after a step-change in the applied voltage (Fig. S3a). It must be noted that our experiments were not optimized for speed but rather aimed at collecting absolute *V*_m_ information from many cells in parallel; recording speed can be improved at the expense of spatial resolution by increasing the frame rate.

Since it is not a priori clear if altered pH has the same impact on τ_lt_ as it has on *F*, we performed a patch-clamp calibration of rEstus-NI in HEK293T cells in extracellular buffers with pH values between 6.4 and 7.9 and found a minor change in τ_lt_ at -120 mV (<50 ps), while *F* relative to the fluorescence intensity of mKate2, which is intracellular, underwent a marked change from 7.5 ± 0.2 at pH 7.9 to 2.0 ± 0.1 at pH 6.4. The median τ_lt_ and the distribution measured for fixed HEK293T cells with a *V*_m_ of approximately 0 mV supports that in the mentioned pH range – given prior calibration τ_lt_ – rEstus-NI provides realistic estimates of *V*_m_ (Fig. S4). It is remarkable how *F* at -120 mV is affected by pH variation compared with the rather pH-resistant fluorescence lifetime (Fig. S4c). However, extracellular changes in pH still requires new calibration of the sensor due to pH-dependent shifts in *V*_h_ and *k*_h_ (Fig. S4d, e). At lower pH, *V*_hF_ and *V*_hτ_ are left-shifted, and *k*_hF_ and *k*_hτ_ become smaller. Both effects are compatible with progressive protonation of the voltage-sensing domain, resulting in an increased effective gating charge (smaller *k*_h_) and a shift in the equilibrium voltage (smaller *V*_h_).

### Analysis of K^+^-channel expressing cells with FLIM measurements

Ideally, FLIM-based *V*_m_ measurements should provide information about individual cells by capturing a FLIM image of many cells. In this proof-of-principle case we studied the impact of overexpressing of large-conductance Ca^2+^- and depolarization-activated K^+^ channels (BK_Ca_, encoded by *KCNMA1*) in HEK293T cells on the resting membrane potential. BK_Ca_ channels are co-activated by an increase in the intracellular Ca^2+^ concentration ([Ca^2+^]_i_) and cell depolarization. Under resting conditions of HEK293T cells with low [Ca^2+^]_i_ and V_rest_ of about -40 mV ^11^, the activity of wild-type BK_Ca_ channels is expected to be close to the detection limit. Therefore, we also examined BK_Ca_ together with its auxiliary subunit BKγ1 (encoded by *LRRC26*), which increases the channel’s activity even under resting conditions ^31^. For current traces in HEK293T cells, see Fig. S6.

Using rEstus-NI in HEK293T cells overexpressing BK_Ca_, we observed that this channel hyperpolarized the cells from a median *V*_m_ of -39.9 mV for mock-transfected HEK293T cells to -48.3 mV (Wilcoxon Rank Test: *p* < 0.001). When the cells were treated with the ionophore gramicidin, their median *V*_m_ changed to -1.4 mV under control conditions and to -11.8 mV for BK_Ca_-containing cells (Fig. 3). In the presence of BKγ1, the cells hyperpolarized even further (median -68.6 mV), and *V*_m_ was shunted with subsequent gramicidin application to -6.1 mV (Fig. 3c,f,h).

**Figure 3.**
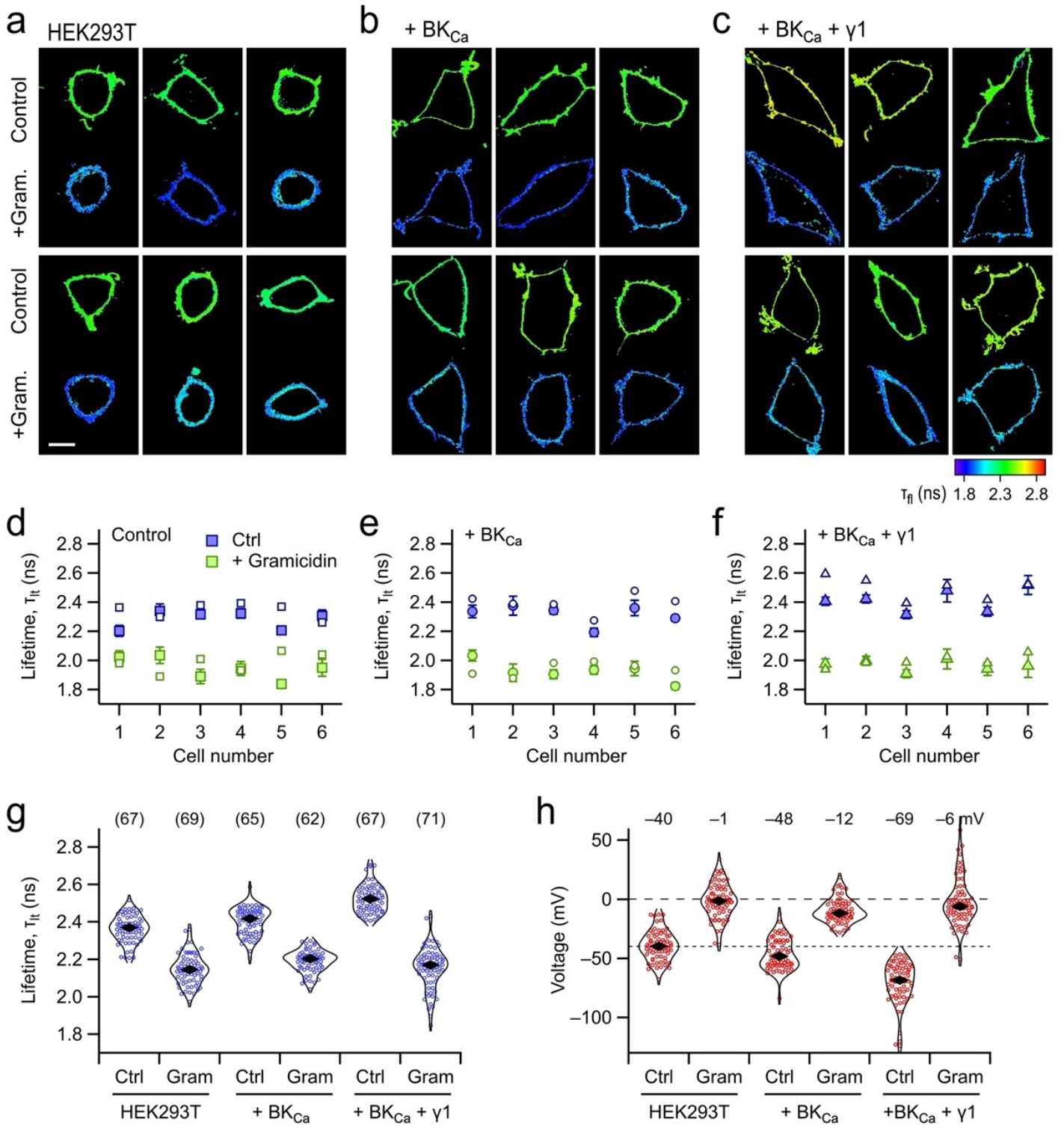
rEstus-NI in HEK293T with BK_Ca_-channel overexpression. (**a**-**c**) FLIM images of HEK293T cells expressing rEstus-NI alone (a), rEstus-NI co-expressed with BK_Ca_ channels (b), and BK_Ca_ α subunits together with the auxiliary subunit BKγ1 (c), without and with treating the cells with gramicidin (1 μM). (**d**-**f**) Lifetime values of HEK293T without (d), and with co-expression of BK_Ca_ channels (e) or BK_Ca_ together with BKγ1 (f) without (blue) and after application of gramicidin (green) for the cells shown in (a-c). The small open symbols are the mean lifetime values determined from the masked mean images shown in (a-c). The filled symbols are median lifetime values of 300 frames with standard deviation from an ROI-based analysis. (**g**) Lifetime values of HEK293T cells without and with overexpression of the indicated K^+^ channels, and without and with gramicidin. The black rhombi indicate the medians of all cells examined, *n* in parentheses. (**h**) The calculated voltages corresponding to each condition, based on the data from (g) and the calibration shown in Fig. S5. The values are the median *V*_m_ estimates. Recordings were performed in extracellular solutions containing 2 mM K^+^ in order to increase the Nernst potential of K^+^ ions; the corresponding calibration in these solutions is shown in Fig. S5.

The estimates of τ_lt_ were homogeneous in the plasma membrane area of individual cells, provided that fluorescence originating from intracellular components is masked. However, there was considerable variability in τ_lt_ between cells (Fig. 3a-f). As expected from the saturation of the τ_fl_(*V*_m_) characteristics at extreme low and high *V*_m_ values (Fig. 2b) and as applicable to all GEVIs with high sensitivity, there are noticeably more outliers after conversion of τ_fl_ to *V*_m_ values used outside its optimum voltage range (Fig. 3h), thus justifying the use of medians for estimating the average *V*_m_ of an ensemble of cells. For rEstus-NI, τ_fl_(*V*) becomes shallow at strong hyperpolarization (≈-100 mV) so that the sensitivity approaches values as determined for ASAP1 (Fig. 2b). However, in its optimum voltage range, the sensitivity of rEstus-NI permitted detecting a slight hyperpolarization of the cells induced by the expression of BK_Ca_ channels (from -40 to -48 mV) when considering the median of >50 cells (Fig. 3h). This result is remarkable because BK_Ca_ channels are not expected to be activated, and hence K^+^ conducting, under resting conditions of *V*_m_ and the intracellular Ca^2+^ concentration of about 100 nM (Fig. S6). It is conceivable that the small hyperpolarization is mediated by occasional BK_Ca_ opening events, triggered by noise in *V*_m_ and/or [Ca^2+^]_i_ of resting HEK293T cells. The activity of BK_Ca_ channels is augmented at resting *V*_m_ and [Ca^2+^]_i_ by coexpression of the BKγ1 subunit (Fig. S6), and this effect was clearly detected in the FLIM measurements as an additional hyperpolarization to about -69 mV. Thus, the combination of BK_Ca_ and rEstus-NI or ASAP3 in HEK293T cells should be suitable for FLIM-based pharmacological screening for BK_Ca_ agonists (so-called BK_Ca_ openers) without the need to manipulate [Ca^2+^]_i_. In the additional presence of BKγ1, also the effect of BK_Ca_ antagonists should be accessible with FLIM by measuring cell depolarization.

### Application of rEstus-NI in mammalian cancer cell lines

The applicability of rEstus-NI to other cell lines was examined by calibrating the sensor in MCF-7 epithelial breast cancer cells (adenocarcinoma) and A375 malignant melanoma cells. The voltage dependencies of *F* and τ_lt_ were similar to the results obtained in HEK293T cells (Fig. 4a-b); for calibration constants see Table S3.

**Figure 4.**
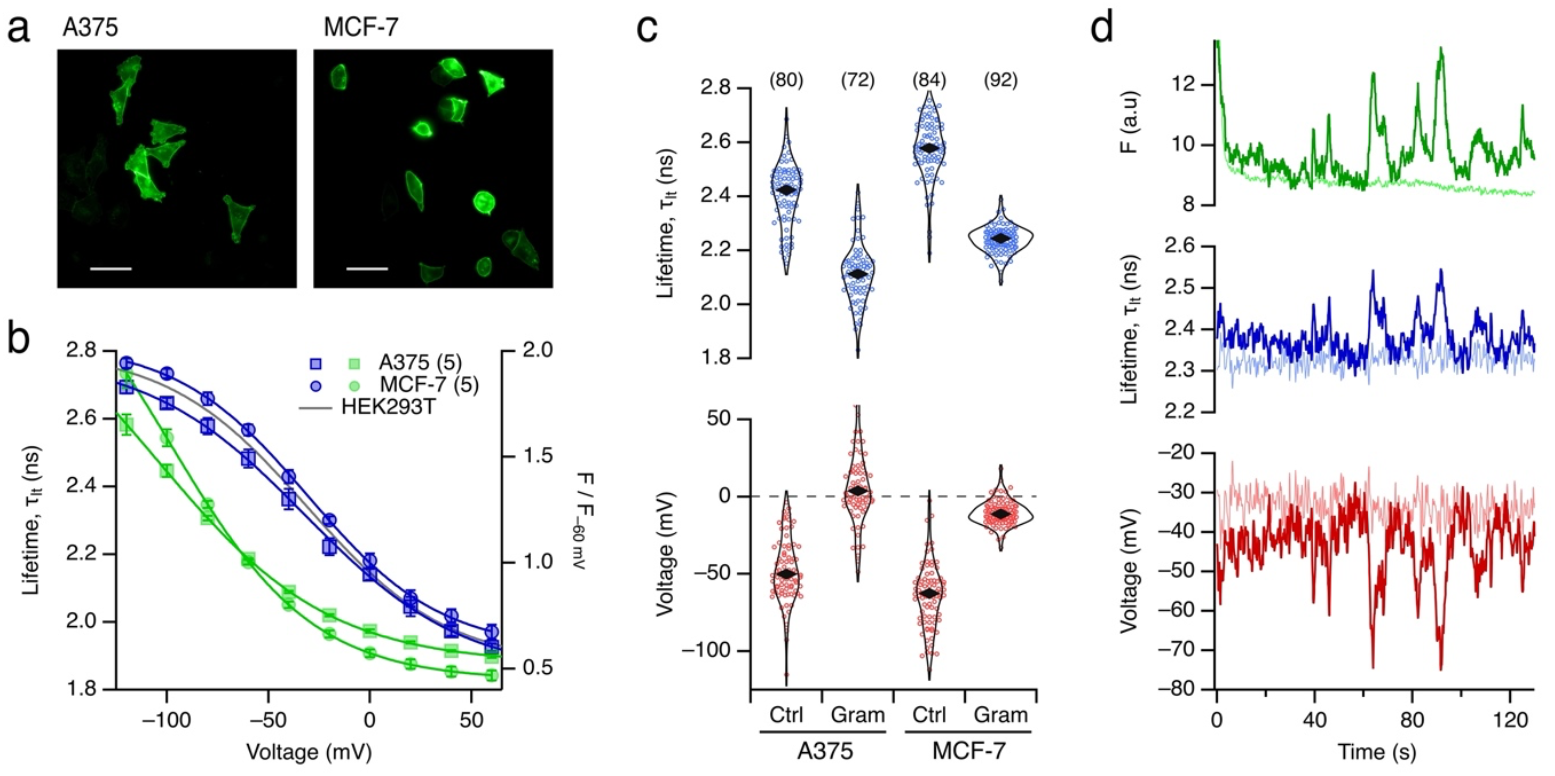
*V*_m_ measurements in cancer cell lines. (**a**) Fluorescence images of A375 melanoma and MCF-7 breast cancer cells, transfected with plasmids coding for rEstus-NI. Scale bars: 50 µm. (**b**) Mean fluorescence intensity, normalized to -60 mV (green), and mean fluorescence lifetime (blue) as a function of voltage for rEstus-NI expressed in A375 and MCF-7 cells, with superimposed fits according to Eq. 1. Error bars indicate ± SEM, and the number of cells measured is given in parentheses. (**c**) *top*: Lifetime values of A375 and MCF-7 cells, without or with 1 μM gramicidin. Each point corresponds to the median lifetime of an individual cell, and the black rhombi indicate the overall median of the cell medians for each condition. *bottom*: The calculated voltages corresponding to each condition, based on the calibration data from (b). (**d**) Time course of *F* (green), _lt_ (blue), and *V*_m_ calculated from _lt_ (red) for A375 cells expressing rEstus-NI, demonstrating that FLIM can detect spontaneous fluctuations in *V*_m_ in these cells. A recording from a cell under voltage-clamp control (−40 mV) is shown as dim lines for comparison to differentiate the intrinsic experimental noise from true voltage fluctuations of A375 cells. For more examples, see Fig. S7.

The distribution of lifetimes in larger cell populations without voltage-clamp control was measured after transient transfection of A375 and MCF-7 cells with plasmid coding for rEstus-NI. Subsequently, the cells were treated with 1 µM gramicidin to short-circuit the cells and, hence, to drive *V*_m_ towards 0 mV. The lifetimes were converted to apparent *V*_m_ values according to the above calibration. From the data shown in Fig. 4c we can derive several conclusions: (i) The violin plots indicate that for A375 and MCF-7 cells there is a wide distribution of apparent *V*_m_ values with obvious extreme outliers. It is therefore appropriate to consider the median of the distributions, which yields a median resting *V*_m_ of -50.2 mV for A375 and -62.7 mV for MCF-7 cells. (ii) After gramicidin application, A375 and MCF-7 cells depolarized as expected, reaching median *V*_m_ values of 3.7 mV and -11.4 mV, respectively (Wilcoxon Rank Test: *p* < 0.001 for both). (iii) Especially for MCF-7 cells the distribution became substantially narrower (standard deviation of 21.7 mV under control conditions reduces to 8.0 mV in gramicidin; Wilcoxon Rank Test for the deviations from the median: *p* < 0.001), suggesting that part of the large *V*_m_ scatter under control conditions must be related to *V*_m_. The same conclusion cannot be drawn directly for A375 cells because the scatter in *V*_m_ was not much affected by gramicidin (*p* = 0.29). If the calibration is based on a representative set of cells, the median *V*_m_ value of a population of cells may still yield realistic estimates, but this estimate does not necessarily apply on a single-cell basis.

We furthermore investigated whether rEstus-NI could detect endogenous *V*_m_ fluctuations in non-excitable cells, potentially by associating true *V*_m_ amplitudes. For this purpose, we recorded the time dependence of *F* and τ_lt_ of individual A375 cells and converted the lifetimes to *V*_m_ estimates. As shown for one such example in Fig. 4d, disregarding the initial drop in fluorescence due to photoswitching, *F* and τ_lt_ fluctuated in a highly correlated manner, with lifetime excursions being on the order of more than 100 ps (blue, median τ_lt_ = 2.37 ns, s.d. = 0.04 ns). In the converted *V*_m_ trace (red), these fluctuations are seen as hyperpolarizing events of about 20-30 mV from a resting voltage of about -40 mV (median *V*_m_ = -42.3 mV, s.d. = 7.8 mV). The events lasted up to about 10 s. For more examples, see Fig. S7.

These spontaneous hyperpolarizing events observed here might be caused by the spontaneous opening of Ca^2+^-activated K^+^ channels (*KCNN4*), which are present in A375 cells ^35^. Recent studies by our group and others have revealed that various non-excitable cells exhibit endogenous electrical activity ^12, 36^. Quicke *et al*. reported spontaneous *V*_m_ fluctuations in various breast cancer cell lines, including the highly aggressive MDA-MB-231 cells ^37^. While the dynamic nature of *V*_m_ in non-excitable cells has only recently gained attention, it is well established that ion channels can contribute to tumor development. For instance, the blockade of *KCNN4*-encoded channels in A375 cells enhances apoptosis ^38^. However, mechanistic insights into the mode of action of ion channels during development and disease progression ^7^ require tools capable of measuring absolute *V*_m_ and its dynamics, and FLIM of ASAP3, rEstus, or rEstus-NI-expressing cells can contribute to this emerging field of science.

## CONCLUSION

Fueled by the demand for non-invasive sensors of the cellular membrane potential, recent developments have resulted in a plethora of fluorescent genetically encoded voltage indicators. Some GEVIs of the ASAP family feature high voltage sensitivity of fluorescence intensity, thus requiring the voltage optimum to be aligned with the sensor application. Moreover, the fluorescence intensity is strongly affected by factors, such as photobleaching, photoswitching, and cell motility. Here we showed how fluorescence lifetime imaging microscopy eliminates most of these confounding factors. Among the ASAP family of GEVIs, ASAP3, rEstus, and rEstus-NI provide a voltage sensitivity of the fluorescence lifetime suitable for measuring absolute *V*_m_ of non-excitable cells. As demonstrated for rEstus-NI, these sensors detect minute alterations of *V*_m_ induced by ion channel activity, are applicable to cancer cell lines, and provide experimental access to the non-invasive recording to membrane voltage dynamics.

## Author Contributions

AGN: generation of expression plasmids and cell lines, electrophysiological and FLIM experiments, data analysis; MR: FLIM experiments, image analysis; PR: generation of expression plasmids, data analysis; JP & MS: infrastructure, FLIM technology, fund raising; TMZ: FLIM experiments, data analysis; SHH: study design, data analysis, fund raising, writing. All authors contributed to manuscript editing.

## Research Funding

Support by the Simons Foundation Autism Research Initiative (SFARI) (SHH, 705944SH), German Academic Exchange Service (AGN, 91819480), and the German Research Foundation, DFG (JP, EXC 2051, Project-ID 390713860). Research infrastructure was supported by the BMBF, funding program Photonics Research Germany (13N15464), the Thüringer Innovationszentrum für Medizintechnik-Lösungen (ThIMEDOP) funded by the European Union within the framework of European Regional Development Fund (ERDF) (IZN 2018 0002), and Freistaat Thüringen / European Funds for Regional Development (EFRE) (2023 HSB 0025). This work is supported by the BMBF, funding program Photonics Research Germany (FKZ: 13N15464.) and is integrated into the Leibniz Center for Photonics in Infection Research (LPI). The LPI initiated by Leibniz-IPHT, Leibniz-HKI, UKJ and FSU Jena is part of the BMBF national roadmap for research infrastructures.

## Data Availability

The data that support the findings of this study are available from the corresponding author upon reasonable request.

## Abbreviations

ASAP: Accelerated sensor of action potentials
BK_Ca_: Large-conductance Ca^2+^- and depolarization-activated K^+^ channel
DMEM: Dulbecco’s modified media
FBS: Fetal bovine serum
FLIM: Fluorescence lifetime imaging microscopy
FRET: Förster resonance energy transfer
GEVI: Genetically encoded voltage indicator
GFP: Green fluorescent protein
PFA: Paraformaldehyde
*V*_m_: Membrane voltage
VSD: Voltage-sensing domain

