## Supplemental Material for "Absolute Membrane Potential Recording with ASAP-Type Genetically Encoded Voltage Indicators Using Fluorescence Lifetime Imaging"

**Supplementary Information**

[a] *Center for Molecular Biomedicine, Department of Biophysics, Friedrich Schiller University Jena and Jena University Hospital, Jena, Germany*

[b] *Leibniz Institute of Photonic Technology, Member of Leibniz Health Technologies, Member of the Leibniz Centre for Photonics in Infection Research (LPI), Albert-Einstein-Straße 9, 07745 Jena, Germany*

[c] *Institute of Physical Chemistry (IPC) and Abbe Center of Photonics (ACP), Friedrich Schiller University Jena, Member of the Leibniz Centre for Photonics in Infection Research (LPI), Helmholtzweg 4, 07743 Jena, Germany, the Cluster of Excellence Balance of the Microverse, Friedrich Schiller University, Jena*

† *Current address: Department of Physics, Polytecnico di Milano, Piazza Leonardo da Vinci 32, 20133 Milan, Italy*

* Corresponding author
*Center for Molecular Biomedicine, Department of Biophysics, Friedrich Schiller University Jena and Jena University Hospital, Hans-Knöll-Straße 2, 07745 Jena, Germany*

ORCID: 0000-0002-4144-0251

### Materials and methods

**Determination of the molecular brightness.** To estimate the molecular brightness of the GEVI constructs (Fig. 2c), they were expressed in HEK293T cells as fusion proteins with mKate2, thus placing mKate2 at the intracellular face of the membrane, while the cpGFP part of the GEVI was located on the extracellular side (Fig. S2a). The voltage of the transiently transfected cells was clamped to -120 mV, and green (GEVI) and red (mKate2) fluorescence were measured photometrically using a Polychrome V monochromator (TILL Photonics, Grafelfing, Germany) and a ProgRes MF cool camera (JenOptik, Jena, Germany). For cpGFP imaging, the excitation wavelength was 480 nm and filter set 38 (Zeiss), in which the excitation filter was replaced with a 492/SP filter (Semrock, Rochester, NY, USA), was used. For mKate2 imaging, the excitation wavelength was 530 nm with filter set 14 (Zeiss). The microscope was an Axio Observer with a 40x EC Plan Neofluar oil-immersion objective (1.3 NA, Zeiss). The relative molecular brightness was defined as the $F_{green}$ / $F_{red}$ ratio at -120 mV.

**Fluorescence Lifetime Image Processing.** To generate fluorescence lifetime images, the second half of the acquired frames were binned using an all-photon filter and individual pixels were fitted with a two-component exponential decay model. This analysis yielded two lifetime images (Tau1, Tau2) and their corresponding intensity images (Int1, Int2). All images were exported as ImageJ-TIFF files with fixed scaling: intensity images were scaled by a factor of 1.0 and lifetime images by 0.001.

An intensity-based thresholding approach was applied to isolate pixels corresponding to the cell membrane while excluding background and noisy regions, without requiring manually drawn ROIs. The threshold was automatically determined by selecting the intensity value that included the top 1000 brightest pixels. A total intensity image (IntSum) was computed by summing int1 and int2, representing the total photon count per pixel. A binary mask was generated by applying a user-defined threshold to IntSum: pixels above the threshold were included and those below were excluded. For each valid pixel, an intensity-weighted average lifetime calculated.

For the analysis of experiments involving many cells, the photon arrival times from these cells (from the second half of the frames, i.e., the last 200 frames out of a total of 400 frames) were binned. A manual ROI was drawn along the cell membrane, and the data were then fitted using a two-component deconvolution fit model.

**Analysis of Time Dependence.** Kinetics in $F$ in response to step-like voltage excursions from -60 mV were described using double-exponential functions for the "on" (after $t_0$) and "off" processes (after $t_1$). The slow time constants for "on" ($\tau_{s,on}$) and "off" ($\tau_{s,off}$) were freely fitted, while the resolution-limited fast time constant ($\tau_f$) and the relative fraction of the slow component ($r_s$) were identical for both the "on" and "off" directions:

$$\frac{F(t<t_0)}{F_{-60mV}} = F_{-60mV}$$ Eq. S2

$$\frac{F(t_0 \le t < t_1)}{F_{-60mV}} = F_{-60mV} + (F_{\max} - F_{-60mV})\left\{\left(1-r_s\right)_1\left(1-\mathrm{e}^{-(t-t_0)/\tau_f}\right) + r_s\left(1-\mathrm{e}^{-(t-t_0)/\tau_{s,on}}\right)\right\}$$

$$\frac{F(t \ge t_1)}{F_{-60mV}} = F_{t_1} - (F_{t_1} - F_{-60mV})\left\{\left(1-r_{off}\right)\left(1-\mathrm{e}^{-(t-t_0)/\tau_f}\right) + \left(1-r_{off}\right)\left(1-\mathrm{e}^{-(t-t_0)/\tau_{s,off}}\right)\right\}$$

with $r_{off} = r_s \dfrac{F_{\max} - F_{-60mV}}{F_{t_1} - F_{-60mV}}$.

**Supplementary Material**

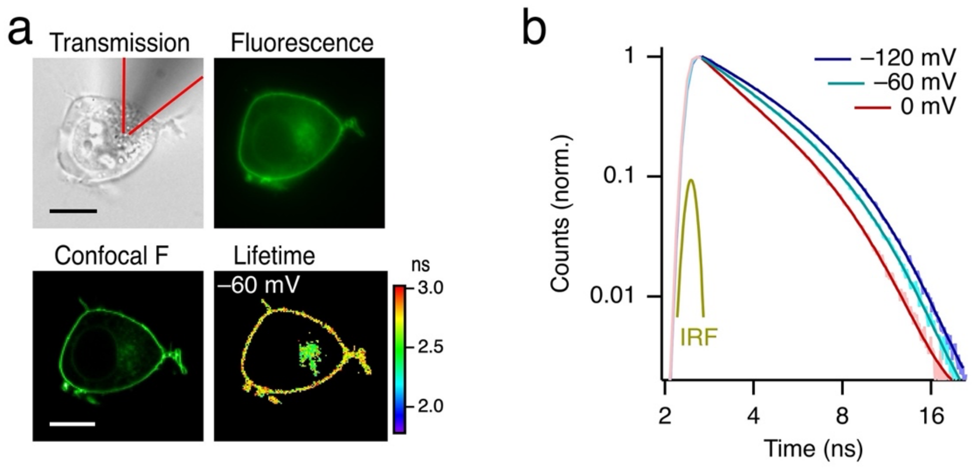


**Supplementary Figure 1. FLIM of HEK293T cells expressing rEstus.** (**a**) Images of a HEK293T cell expressing rEstus with a patch-clamp pipette attached: (i) transmission image with the red lines indicating the patch-clamp pipette, which is out of focus; (ii) raw fluorescence with excitation at 488 nm; (iii) confocal fluorescence intensity of the membrane; (iv) mean intensity-weighted fluorescence lifetime of 20 consecutive images similar to (iii) after masking pixels with too small intensity. Scale bar: 10 µm. (**b**) Photon counts (normalized to the maximum) originating from a membrane-delimiting ROI, as a function of time in response to the indicated internal response function (IRF) for the specified voltages. Thin traces are raw data, and thick lines are double-exponential fits after deconvolution of the IRF.

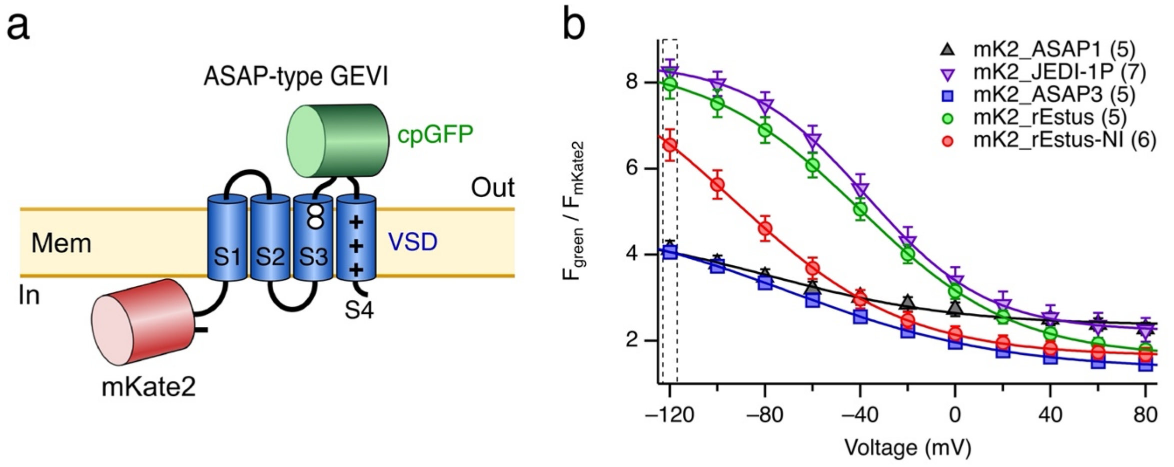


**Supplementary Figure 2. mKate2-GEVI fusion for brightness determination.** (**a**) Scheme of the mKate2-GEVI fusion constructs used for brightness calibration. The GEVIs of the ASAP family have in common a voltage-sensing domain (VSD) consisting of four transmembrane helices (S1-S4, blue) and a circularly permuted GFP (cpGFP) inserted on the extracellular side between S3 and S4. The white circles in S3 mark the approximate positions of residues G138 and T141 of rEstus. (**b**) Voltage dependence of the fluorescence ratio $F_{green}$ / $F_{mKate2}$, originating from the cpGFP moiety of the voltage-dependent GEVIs and the N-terminally fused mKate2, respectively. The mKate2 fluorescence serves as a calibration signal to compare the brightness of the indicated GEVIs at -120 mV (see Fig. 2c). Data are means ± SEM, with *n* in parentheses. Superimposed curves are fit results according to Eq. (1); the resulting parameters are listed in Table S1.

**Supplementary Table 1. Voltage dependence of GEVI fluorescence relative to mKate2 fluorescence.** Results of the data fits as shown in Fig. S2 according to Eq. (1). Note that the fit for ASAP1 was constrained regarding to a reasonable steepness ($k_h$ = 40 mV) because, even at -120 mV, the fluorescence is far from being saturated. Errors are 95% confidence intervals.

| mKate2 fusion with | Max. $F_{green}$ / $F_{red}$ | Offset (%) | $V_{hF}$ (mV) | $k_{hF}$ (mV) | n |
|---|---|---|---|---|---|
| ASAP1 | 4.66 ± 0.19 | 50.6 ± 1.9 | -78.3 ± 8.9 | 40, constr. | 5 |
| JEDI-1P | 8.48 ± 0.29 | 26.3 ± 0.3 | -37.2 ± 0.4 | 25.4 ± 0.4 | 7 |
| ASAP3 | 4.96 ± 0.02 | 26.8 ± 0.2 | -69.9±0.6 | 45.0 ± 0.4 | 5 |
| rEstus | 8.49 ± 0.04 | 19.1 ± 0.3 | -40.3 ± 0.4 | 32.6 ± 0.5 | 5 |
| rEstus-NI | 8.87 ± 0.16 | 18.5 ± 0.1 | -92.9 ± 1.7 | 35.8 ± 0.7 | 6 |

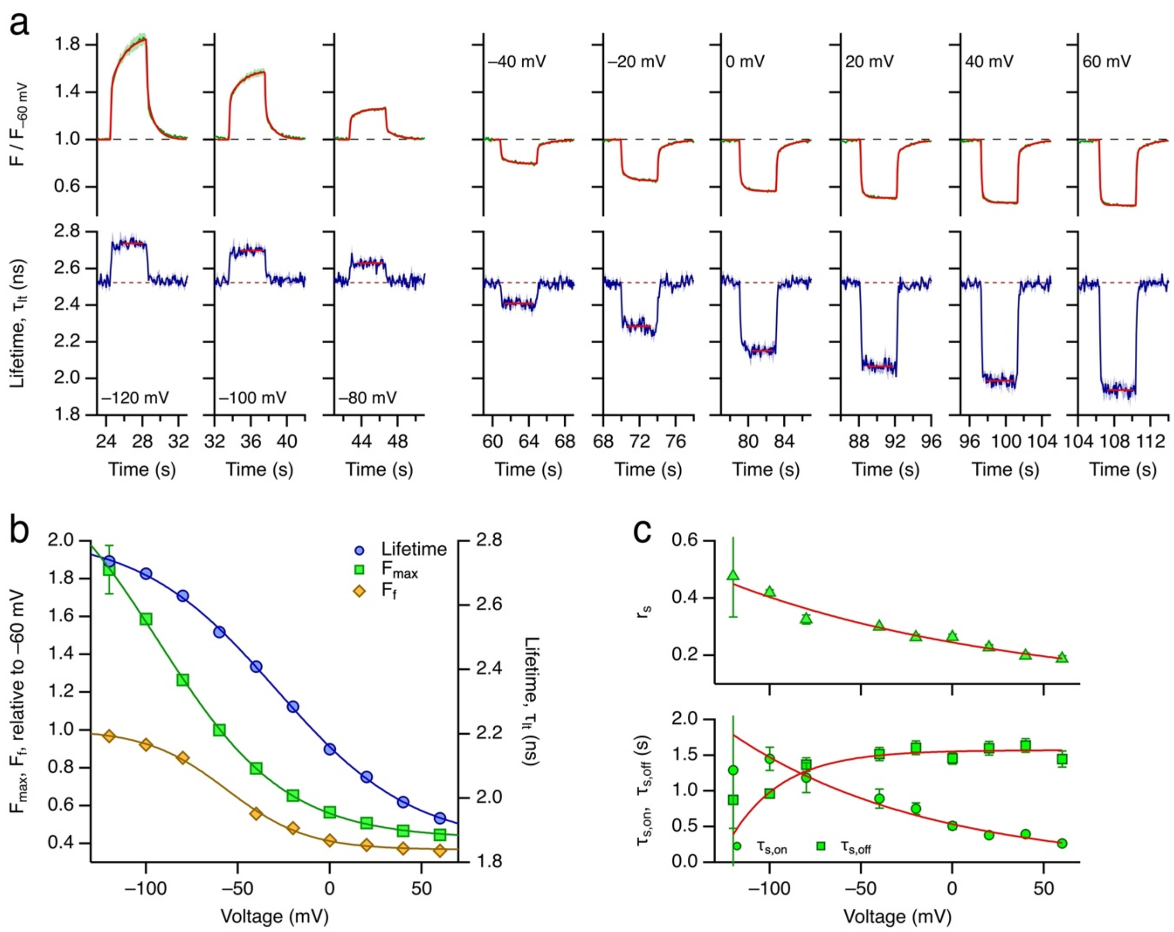


**Supplementary Figure 3. Kinetics of voltage-step responses in fluorescence and fluorescence lifetime of rEstus-NI.** (**a**) Responses of *F* normalized to *F* at -60 mV (*top*) and fluorescence lifetime ($\tau_{lt}$) (*bottom*) to voltage steps to the indicated voltage from and to -60 mV (data from Fig. 1e). The *F* data are superimposed with fit functions according to Eq. (S2), thus comprising double-exponential "on" and "off" phases with a common fast time constant ($\tau_f$, 70 ms) and a common fraction of the fast component ($r_s$). The red lines superimposed on the lifetime data show the data range used for determining the mean. (**b**) Voltage dependence of the maximal fluorescence intensity ($F_{max}$), the intensity of the fast component only ($F_f$), and the fluorescence lifetime with superimposed fits according to Eq. (1). For the resulting parameters, see Table S2. (**c**) Results of the kinetics analysis showing the voltage dependence of the fraction of the slow component ($r_s$, *top*), and the time constants of the slow component for the voltage step from ($\tau_{s,on}$) and the return to -60 mV ($\tau_{s,off}$). The superimposed curves are fitted single-exponential functions to indicate the trend of voltage dependence.

**Supplementary Table 2. Voltage dependence of rEstus-NI fluorescence components based on double-exponential fits according to Eq. (2).** The table contains the parameters describing the data fits according to Eq. (1) shown in Fig. S3b.

| Parameter | $\Delta a$ | $a_{inf}$ | $V_h$ (mV) | $k_h$ (mV) |
|---|---|---|---|---|
| Lifetime, $\tau_{lt}$ | 0.96 ± 0.03 ns | 1.86 ± 0.02 ns | -29.2 ± 1.4 | 37.3 ± 1.9 |
| $F_{max}$ | 2.09 ± 0.09 | 0.43 ± 0.01 | -93.4 ± 2.9 | 34.8 ± 1.4 |
| $F_f$ | 0.63 ± 0.02 | 0.37 ± 0.01 | -56.0 ± 2.3 | 21.9 ± 1.7 |

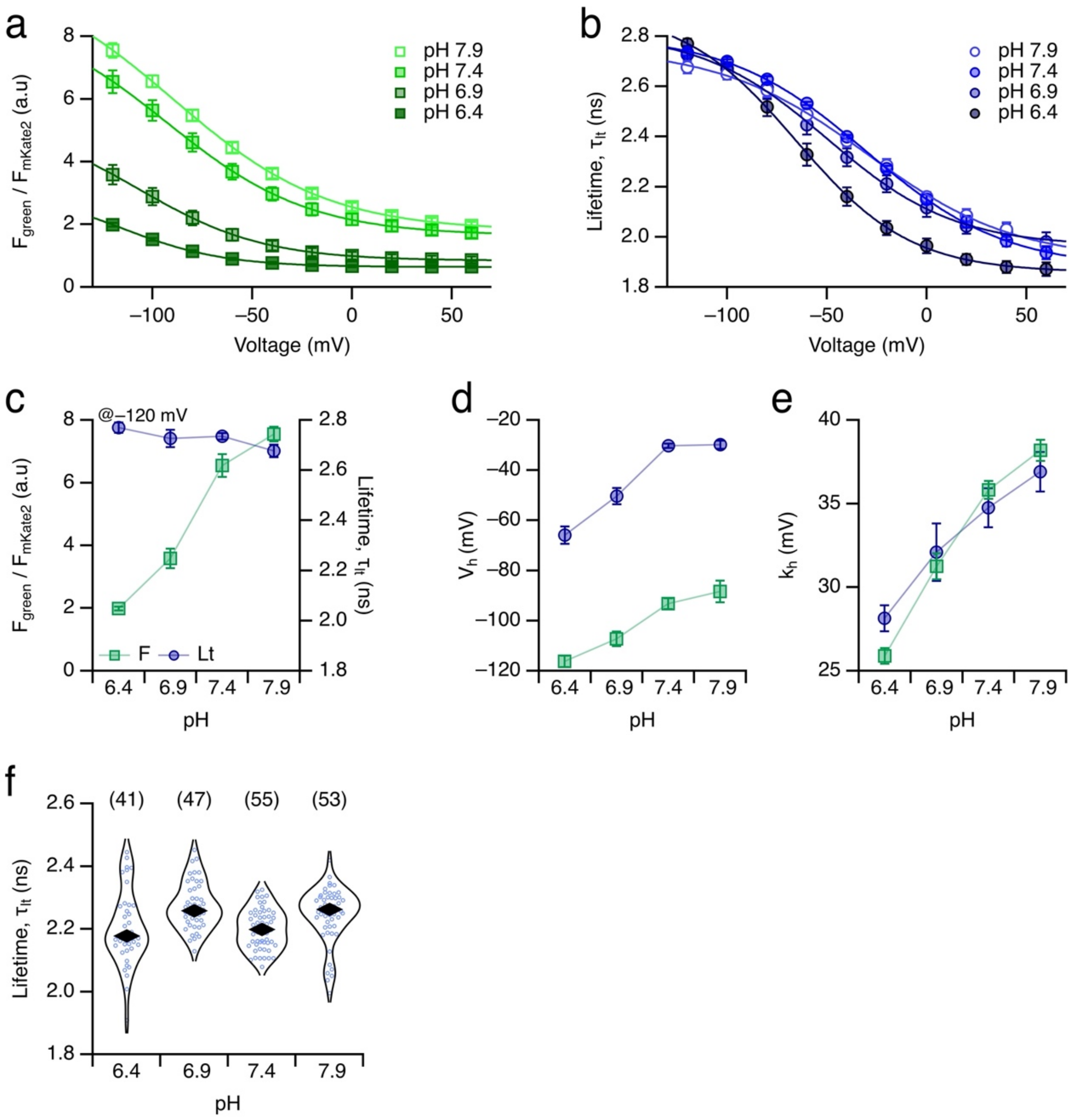


**Supplementary Figure 4. Dependence of rEstus-NI on extracellular pH.** rEstus-NI was expressed in HEK293T cells. (**a**) Molecular brightness relative to the fluorescence of mKate2 (as in Fig. S2) as a function of voltage for the indicated extracellular pH values. The intracellular (pipette) solution had pH 7.4 in all cases. Data are means ± SEM for *n* = 5 cells. The superimposed curves are data fits according to Eq. (1); the resulting parameters are listed in Table S3. (**b**) Fluorescence lifetime as a function of voltage of the same cells as in (a). (**c**-**e**) Parameters derived from the data shown in (a) and (b) as a function of pH: Relative brightness and fluorescence lifetime at -120 mV (**c**), half-maximal voltages, $V_h$ (**d**), and the slope factors, $k_h$ (**e**) for the fluorescence intensity (squares) and the fluorescence lifetime (circles). (**f**) Distributions of fluorescence lifetime values for fixed HEK293T cells expressing rEstus-NI in extracellular buffers with the indicated pH values. Each blue point corresponds to the median lifetime of an individual cell, and the black rhombi indicate the overall median of the cell medians for each condition. Cell fixation not only depolarizes the cells but also permeabilizes the membrane such that the indicated pH values also apply to the intracellular side.

**Supplementary Table 4. Voltage dependence of rEstus-NI fluorescence and fluorescence lifetime in HEK293T cells at different extracellular pH.** The table contains the parameters describing the data fits shown in Fig. S4. Errors are 95% confidence intervals.

| pH | Parameter | $\Delta a$ | $a_{inf}$ | $V_h$ (mV) | $k_h$ (mV) | n |
|---|---|---|---|---|---|---|
| 7.9 | $F_{green}$ / $F_{mKate2}$ | 8.38 ± 0.09 | 1.7 ± 0.01 | -89.3 ± 0.8 | 38.7 ± 0.4 | 5 |
| | Lifetime, $\tau_{lt}$ | 0.85 ± 0.04 ns | 1.91 ± 0.02 ns | -29.7 ± 2.1 | 37.1 ± 2.9 | 5 |
| 7.4 | $F_{green}$ / $F_{mKate2}$ | 7.23 ± 0.17 | 1.64 ± 0.01 | -92.9 ± 1.7 | 35.8 ± 0.7 | 5 |
| | Lifetime, $\tau_{lt}$ | 0.93 ± 0.02 ns | 1.88 ± 0.01 ns | -30.2 ± 0.9 | 34.8 ± 1.2 | 5 |
| 6.9 | $F_{green}$ / $F_{mKate2}$ | 4.54 ± 0.12 | 0.84 ± 0.01 | -106.6 ± 1.7 | 31.3 ± 0.6 | 5 |
| | Lifetime, $\tau_{lt}$ | 0.85 ± 0.04 ns | 1.96 ± 0.01 ns | -49.7 ± 2.3 | 31.9 ± 2.4 | 5 |
| 6.4 | $F_{green}$ / $F_{mKate2}$ | 2.51 ± 0.03 | 0.64 ± 0.01 | -116.1 ± 0.6 | 26.0 ± 0.2 | 5 |
| | Lifetime, $\tau_{lt}$ | 1.05 ± 0.09 ns | 1.86 ± 0.02 ns | -65.4 ± 5.6 | 28.7 ± 4.3 | 5 |

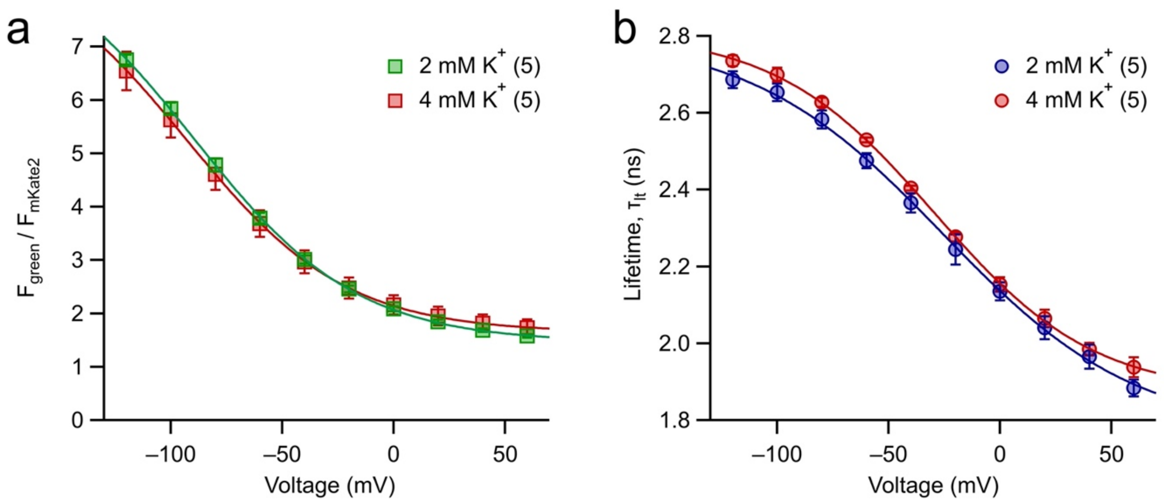


**Supplementary Figure 5. Calibration of rEstus-NI in 2-mM $K^+$ buffer.** rEstus-NI was expressed in HEK293T cells and fluorescence intensity (**a**) and fluorescence lifetime (**b**) were measured under voltage-clamp control in the whole-cell configuration. Extracellular solutions either contained 4 or 2 mM $K^+$, as indicated. The superimposed curves are results of data fits according to Eq. (1), yielding the calibration parameters presented in Table S4.

**Supplementary Table 4. Voltage dependence of rEstus-NI fluorescence and fluorescence lifetime in HEK293T cells at different extracellular $K^+$ concentrations.** The table contains the parameters describing the data fits shown in Fig. S5. Errors are 95% confidence intervals.

| $[K^+]_o$ (mM) | Parameter | $\Delta a$ | $a_{inf}$ | $V_h$ (mV) | $k_h$ (mV) | n |
|---|---|---|---|---|---|---|
| 2 | $F_{green}$ / $F_{mKate2}$ | 7.60 ± 0.20 | 1.46 ± 0.02 | -89.3 ± 1.8 | 36.5 ± 0.9 | 5 |
| | Lifetime, $\tau_{lt}$ | 1.04 ± 0.06 ns | 1.77 ± 0.03 ns | -25.7 ± 2.6 | 43.2 ± 3.8 | 5 |
| 4 | $F_{green}$ / $F_{mKate2}$ | 7.23 ± 0.17 | 1.64 ± 0.01 | -92.9 ± 1.7 | 35.8 ± 0.7 | 5 |
| | Lifetime, $\tau_{lt}$ | 0.94 ± 0.02 ns | 1.87 ± 0.01 ns | -29.4 ± 1.7 | 35.8 ± 0.7 | 5 |

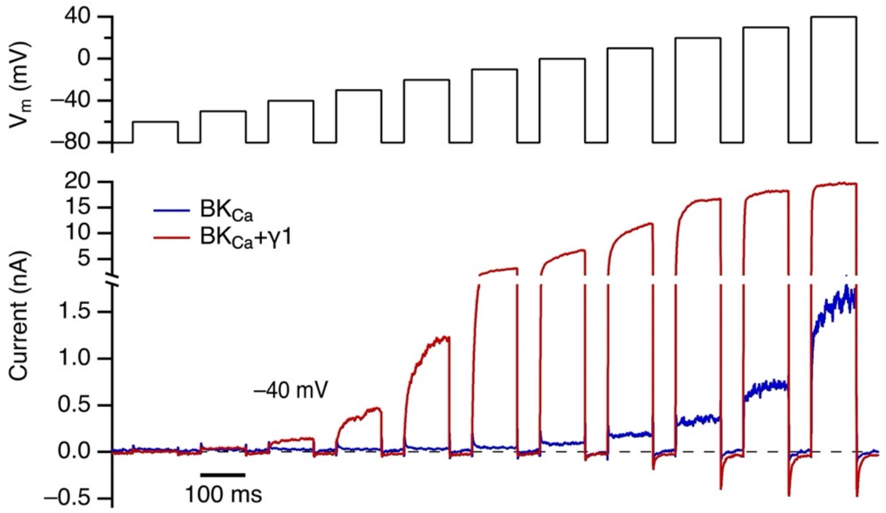


**Supplementary Figure 6. Function of $BK_{Ca}$ and $BK_{Ca}$/BKγ1 channels in HEK293T cells.** Whole-cell current recordings according to the pulse protocol shown in the top panel. HEK293T cell expressed rEstus-NI along with $BK_{Ca}$ (Slo1, *KCNMA1*, blue) or $BK_{Ca}$ together with BKγ1 (*KCNMA1-LRRC26*, red). Traces are means of three recordings. In the presence of BKγ1 current saturated at around 0 mV. The intracellular solution contained 100 nM free $Ca^{2+}$. Under these conditions, channel activity of $BK_{Ca}$ alone is very low at -40 mV whereas in the presence of BKγ1 outward $K^+$ current is clearly discernable. Pipette solution in mM: 135 KCl, 2 $MgCl_2$, 3.93 $CaCl_2$, 10 EGTA, 10 HEPES, pH 7.4 (KOH).

**Supplementary Table 5. Voltage dependence of rEstus-NI fluorescence and fluorescence lifetime in human cancer cell lines.** The table contains the parameters describing the data fits shown in Fig. 4b for A375 and MCF-7 cells. Errors are 95% confidence intervals. Data for HEK293T cells are given as reference. In addition to the lifetime calibration based on the averages of lifetime frames, we also provide a calibration based on the medians to minimize the influence of extreme outliers.

| Cell Type | Parameter | $a_{max}$ | $a_{inf}$ | $V_h$ (mV) | $k_h$ (mV) | n |
|---|---|---|---|---|---|---|
| A375 | *F* | 2.44 ± 0.07 | 1.92 ± 0.01 | -104.6 ± 3.0 | 42.4 ± 1.2 | 5 |
| | Lifetime, $\tau_{lt}$ | 2.79 ± 0.02 ns | 0.89 ± 0.03 ns | -36.8 ± 1.8 | 38.6 ± 2.4 | 5 |
| median | Lifetime, $\tau_{lt}$ | 2.79 ± 0.05 ns | 0.94 ± 0.08 ns | -32.0 ± 4.5 | 39.0 ± 5.9 | 5 |
| MCF-7 | *F* | 1.98 ± 0.07 | 1.53 ± 0.08 | -79.4 ± 3.0 | 41.4 ± 1.2 | 5 |
| | Lifetime, $\tau_{lt}$ | 2.76 ± 0.01 ns | 0.85 ± 0.03 ns | -24.0 ± 1.6 | 40.1 ± 2.2 | 5 |
| median | Lifetime, $\tau_{lt}$ | 2.74 ± 0.13 ns | 0.84 ± 0.24 ns | -29.9 ± 13.1 | 40.8 ± 19 | 5 |
| HEK293T | *F* | 2.52 ± 0.09 | 2.09 ± 0.09 | -93.4 ± 2.9 | 34.8 ± 1.4 | 5 |
| | Lifetime, $\tau_{lt}$ | 2.82 ± 0.02 ns | 0.96 ± 0.03 ns | -29.2 ± 1.4 | 37.3 ± 1.9 | 5 |
| median | Lifetime, $\tau_{lt}$ | 2.81 ± 0.03 ns | 0.93 ± 0.06 ns | -30.8 ± 2.7 | 34.7 ± 3.5 | 5 |

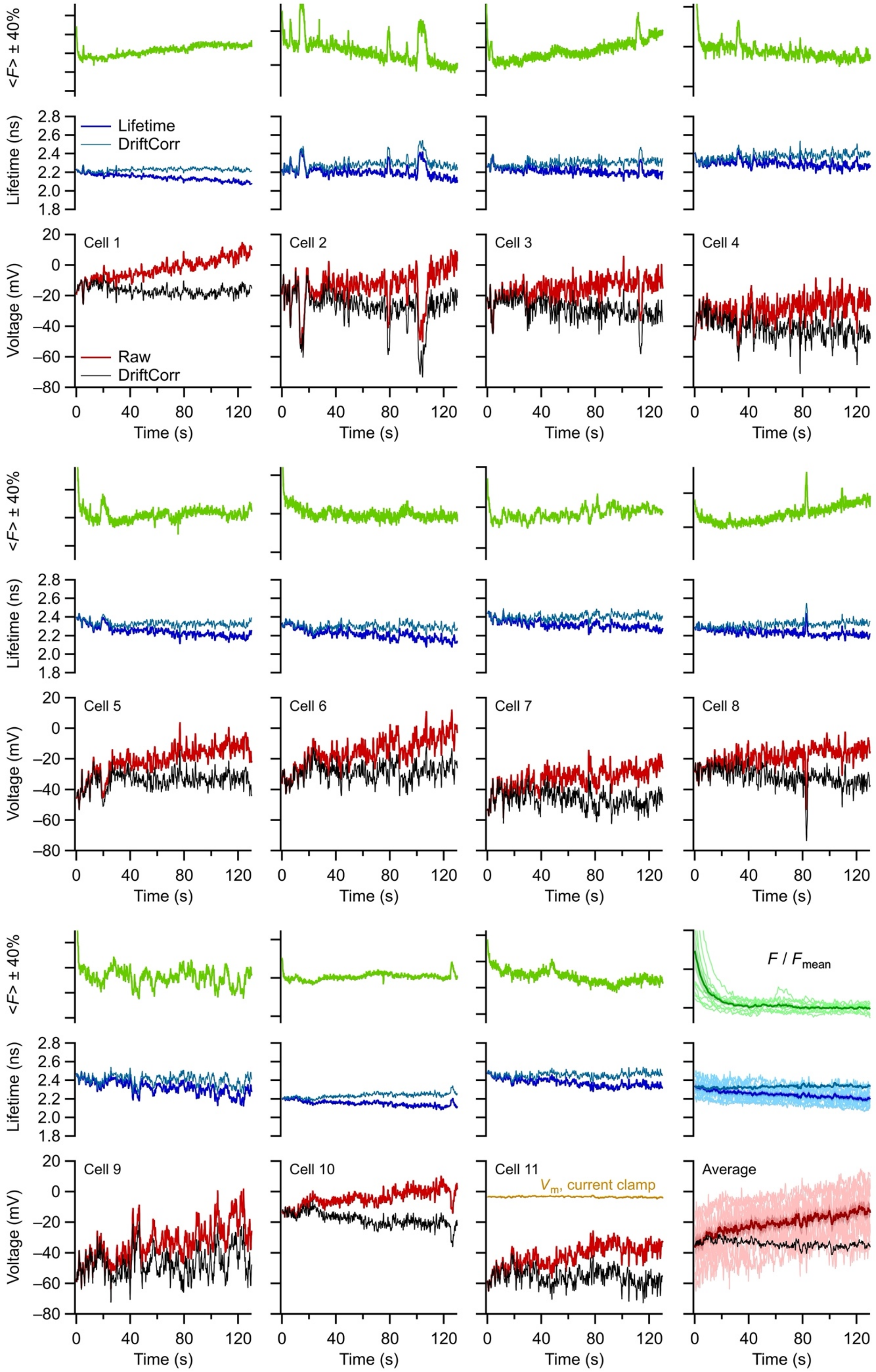

<F> ± 40%
Lifetime (ns)
Lifetime
DriftCorr
Voltage (mV)
Raw
DriftCorr
Cell 1
Cell 2
Cell 3
Cell 4
Cell 5
Cell 6
Cell 7
Cell 8
Cell 9
Cell 10
Cell 11
Average
$V_m$, current clamp
$F$ / $F_{mean}$
Time (s)

**Supplementary Figure 7. Spontaneous $V_m$ fluctuations in A375 cells.** Time courses of FLIM recordings from individual rEstus-NI-transfected A375 cells with the fluorescence intensity (*top*) as mean fluorescence ± 40%, the fluorescence lifetime (*center*), and the voltage derived from the lifetime (*bottom*). The last panels (*bottom right*) display all traces (thin lines) with superimposed mean (thick line) and SEM in shading. Raw lifetime traces (dark blue) are also presented after drift correction (light blue) based on the mean drift in $\tau_{lt}$ measured under whole-cell voltage clamp at -40 mV. Similarly, the resulting voltage is shown based on raw lifetime traces (dark red) and on drift-corrected lifetimes (black). In the panels of the averaged traces, only the mean drift-corrected lifetime and voltage are presented in addition to the individual raw data. The orange trace in the voltage panel of Cell 11 is a voltage recording in current-clamp mode to illustrate the noise level when $V_m$ is measured electrically with intracellular $Ca^{2+}$ chelated with 10 mM EGTA.
